# Genetic architecture constrains exploitation of siderophore cooperation in *Burkholderia cenocepacia*

**DOI:** 10.1101/583898

**Authors:** Santosh Sathe, Anugraha Mathew, Kirsty Agnoli, Leo Eberl, Rolf Kümmerli

## Abstract

Explaining how cooperation can persist in the presence of cheaters, exploiting the cooperative acts, is a challenge for evolutionary biology. While microbial systems have proved extremely useful to test evolutionary theory and identify mechanisms maintaining cooperation, our knowledge is often limited to insights gained from a few model organisms. Here, we introduce siderophore secretion by the bacterium *Burkholderia cenocepacia* as a novel study system. Using a combination of phenotypic and competition assays we found that ornibactin, the main siderophore used for iron scavenging in this species, is secreted into the media, can be shared as public good between cells, but cannot be exploited by ornibactin-defective mutants. Molecular analysis revealed that cheating is compromised because the ornibactin receptor gene and genes involved in ornibactin synthesis are co-expressed from the same operon, such that disruptive mutations in the upstream synthesis genes compromise receptor availability. To prove that it is the genetic architecture of the siderophore locus that prevents cheating, we broke the linked traits by expressing the ornibactin receptor from a plasmid, a measure that turned the ornibactin mutant into a functional cheater. A literature survey across *Burkholderia* species suggests that the genetic linkage independently broke over evolutionary time scales in several lineages, indicating that cheating and countermeasures might be under selection. Altogether, our results highlight that expanding our repertoire of microbial study systems leads to new discoveries and reinforce the view that social interactions shape evolutionary dynamics in microbial communities.

---

Siderophores are secondary metabolites secreted by bacteria to scavenge insoluble and host-bound iron from the environment^1,2^. Siderophores are of major ecological importance as they fulfil a wide range of functions. They constitute virulence factors in infections^1,3^; remediate heavy-metal polluted environments^4,5^; and drive community dynamics by supressing the growth of competitors^6^ or benefiting close relatives through cooperative molecule sharing^7,8^. Especially the observation that the secretion of siderophores can constitute a cooperative act, benefiting individuals other than the producers, has attracted enormous interdisciplinary attention^7,9–14^. The key question in this context is how cooperative siderophore secretion can be evolutionarily maintained, given that siderophore-negative cheater mutants can arise and freeride on the public goods produced by others^15,16^. A large body of work has tackled this question and revealed that cheating and cheater control are major determinants of bacterial population dynamics in infections and in laboratory and natural communities^6,9,11,17–19^. However, despite the great advance these insights provide for our understanding of microbial social interactions and community dynamics, most of the work carried out so far, with a few notable exceptions^9,20,21^, stem from one type of siderophore (pyoverdine) produced by one type of bacterium (fluorescent pseudomonads, particularly *Pseudomonas aeruginosa*).

This limitation prompted us to test whether the findings reported for *P. aeruginosa* are generalizable and applicable to other bacterial systems. We thus set out to explore the social role of the two siderophores, ornibactin and pyochelin, produced by *Burkholderia cenocepacia*^22–24^ This bacterium is, like *P. aeruginosa*, an opportunistic pathogen that can inhabit a wide range of environments^25,26^. Using this study system, we repeated a number of key experiments, which previously demonstrated the social nature of pyoverdine in *P. aeruginosa*^27–30^. Specifically, we carried out siderophore secretion and supernatant cross-feeding assays, and competed a siderophore-negative mutant against producer strains across a range of conditions by manipulating competition time, culture mixing, strain density and frequency. While our results revealed that both siderophores of *B. cenocepacia* are secreted into the environment, we consistently found that only pyochelin but not ornibactin is an exploitable public good in this species.

This surprising finding motivated us to investigate the molecular basis of why ornibactin cannot be exploited in *B. cenocepacia*. We suspected that the genetic architecture of the siderophore locus could play a key role in determining whether non-producers can exploit a specific siderophore. Specifically, we predict that cheating relies on the independent regulation of siderophore synthesis and receptor genes, which enables a strain deficient in siderophore production to express the receptor required for siderophore uptake. Such an independent regulation is indeed in place for pyoverdine in *P. aeruginosa*^31^. For *B. cenocepacia*, the literature tells us that genes encoding the pyochelin synthesis machinery and the receptor are organised in different operons^32^ and might thus also be regulatorily independent. Conversely, the ornibactin receptor gene is located downstream of the synthesis genes as part of the same operon^22,32^, indicating strong positive regulatory linkage between siderophore synthesis and uptake. Here, we use a combination of gene expression analysis and strain engineering to test whether a deletion in a synthesis gene, which turns a producer into a non-producer, negatively affect the expression of downstream receptor genes, and thus compromise the evolutionary success of non-producers.

## Results

### *B. cenocepacia* H111 secretes ornibactin and pyochelin into the media

We used *B. cenocepacia* H111 and its isogenic siderophore mutants^23^ for all our experiments. We first used the colorimetric CAS assay to confirm that the siderophores ornibactin and pyochelin are produced and secreted into the extracellular growth medium under iron limitation in casamino acid (CAA) medium (Fig. 1a). We indeed observed significant CAS activities in all producer strains, although activities differed between them (ANOVA: *F*_3,24_ = 3482; *p* < 0.0001, Fig. 1b). CAS activity was highest for the wildtype supernatant followed by a stepwise decline from H111Δ*pchAB* (ornibactin producer) to H111Δ*orbJ* (pyochelin producer). The higher CAS activity for H111Δ*pchAB* is probably due to ornibactin having a higher iron affinity than pyochelin^33^. As expected, the supernatant of the siderophore-deficient strain H111Δ*orbJ*Δ*pchAB* showed almost completely abrogated CAS activity. Altogether, these results demonstrate that H111 secretes siderophores into the media, which can potentially be shared as public good among cells.

**Figure 1.**
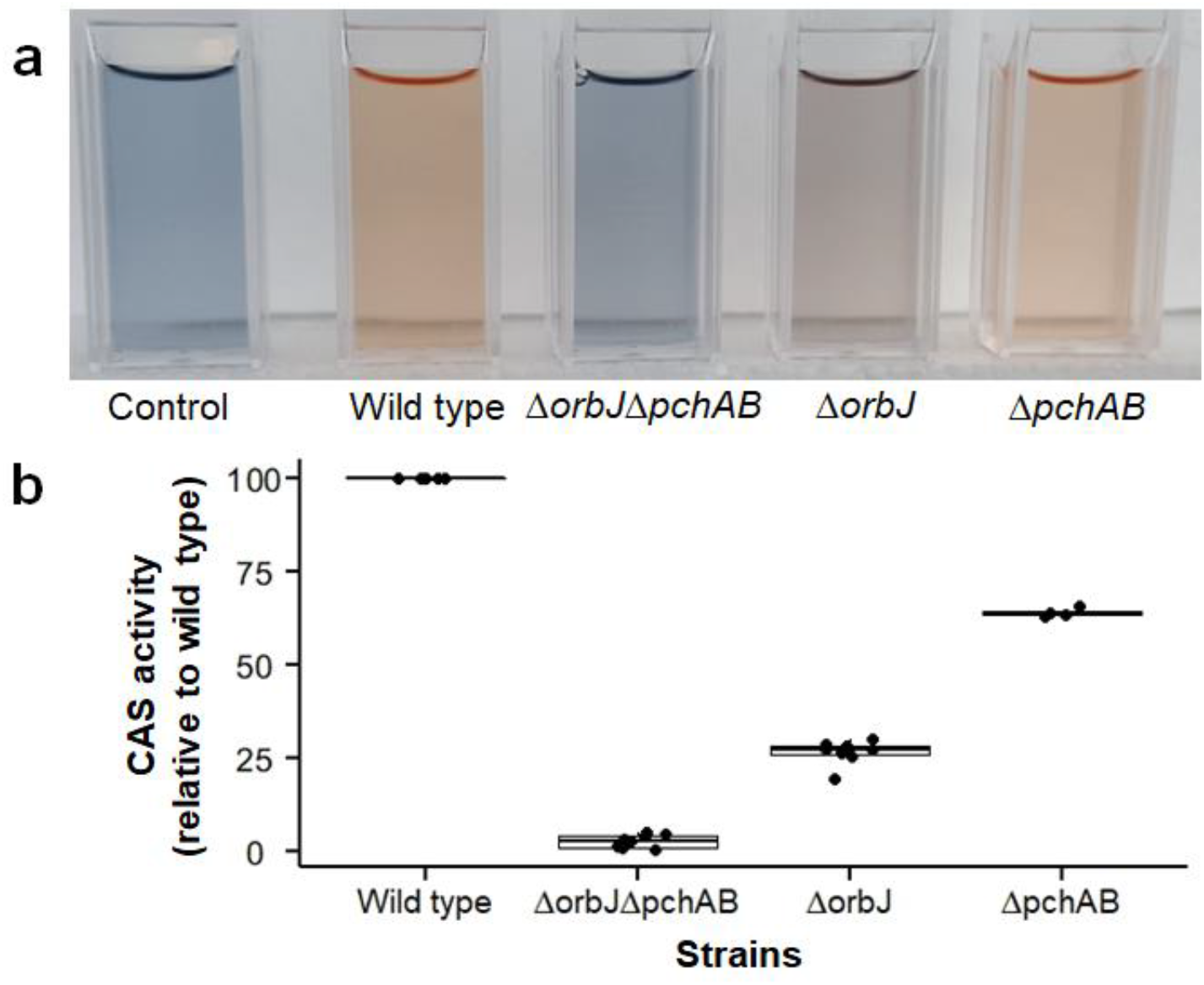
*B. cenocepacia* H111 secretes ornibactin and pyochelin into the media. **a**, the colorimetric CAS assay detects siderophore activity in the extracellular cell-free supernatant by inducing a color change from blue to orange. b, the quantification of this color change through absorbance at 630 nm shows no siderophore activity in the CAA medium and for the double mutant (H111Δ*orbJ*Δ*pchAB*), which is unable to produce ornibactin and pyochelin. The supernatant of the wild type H111 showed highest siderophore activity, followed by the ornibactin producer (H111ΔpchAB) and the pyochelin producer (H111Δ*orbJ*).

### Non-producers can freely use secreted pyochelin whereas access to ornibactin is restricted

We collected the supernatants from siderophore-producing strains to test whether the siderophore non-producer (H111Δ*orbJ*Δ*pchAB*) can use the secreted siderophores for growth. We found that the growth of H111Δ*orbJ*Δ*pchAB* was stimulated in supernatants containing siderophores (Fig. 2a), demonstrating that siderophores can be shared between cells. However, there were significant donor effects with regard to the level of growth stimulation (F_4,27_ = 96.5; *p* < 0.0001). While non-producer cells grew equally well in the supernatants from the wild type and H111Δ*orbJ*, stimulation was significantly compromised when grown in the supernatant of H111Δ*pchAB* (LM: *t*_27_ = 8.76; *p* < 0.0001). This result was a first indication that ornibactin might be less accessible to non-producers than pyochelin. Our control experiments confirmed that the observed growth stimulation was directly driven by siderophores, because all effects disappeared when H111Δ*orbJ*Δ*pchAB* grew with supernatants without siderophores (Fig. 2b, *F*_3,28_ = 2.56; *p* = 0.0748) or in iron-rich medium, where siderophores are not required (Fig. S1).

**Figure 2.**
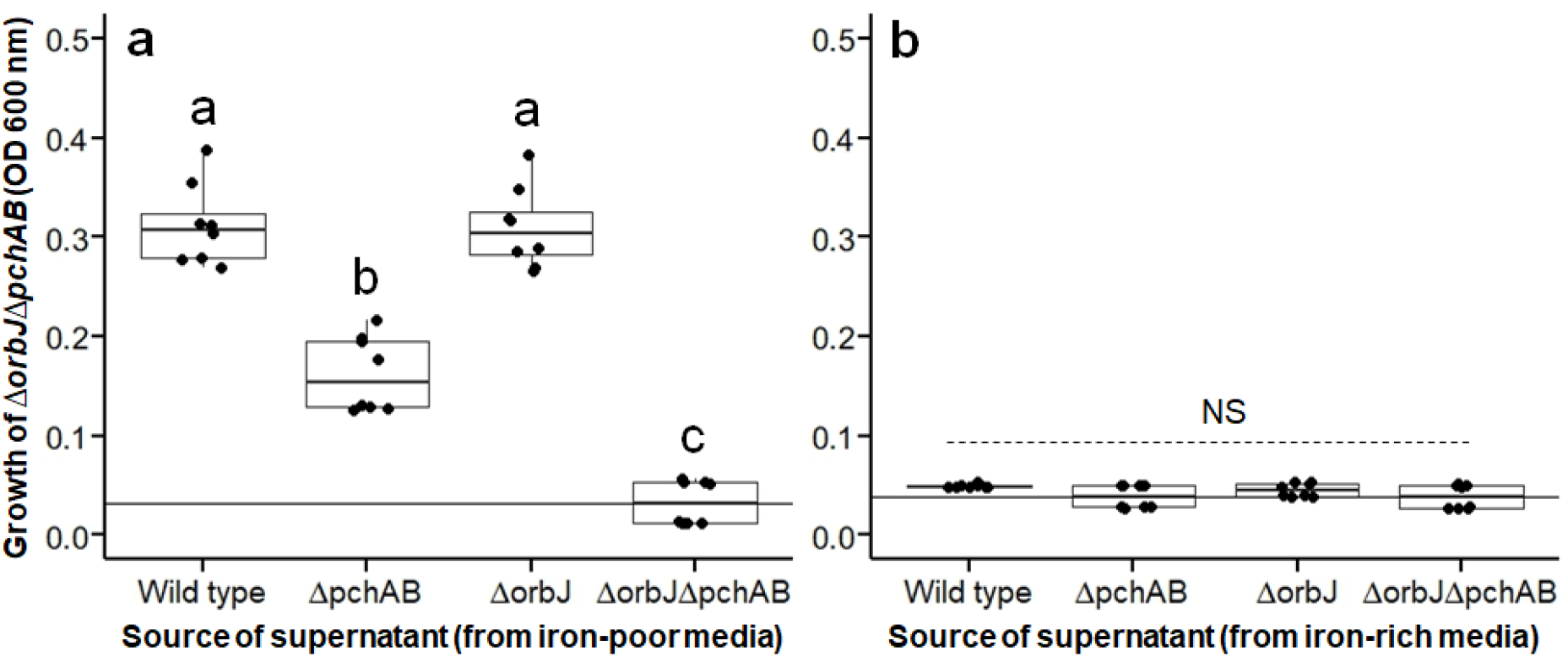
The siderophore non-producer H111Δ*orbJ*Δ*pchAB* can take up supplemented siderophores, but growth induction differs between siderophore types. We tested whether the siderophore non-producer H111Δ*orbJ*Δ*pchAB* can benefit from exogenous siderophores present in the supernatant of donor strains. **a**, supernatants from siderophore producers grown for 24 hours in iron-poor media significantly stimulated the growth of the siderophore non-producer H111Δ*orbJ*Δ*pchAB*. However, growth stimulation was reduced with supernatants from the H111Δ*pchAB* donor strain containing only ornibactin. **b**, in our control experiments, the growth-stimulatory effects disappeared when supernatants were harvested from iron-rich media, which contain little or no siderophores. Different letters above the boxplots indicate statistically significant differences between treatments.

### Competition assays suggest that only pyochelin but not ornibactin is an exploitable siderophore

An important feature of microbial cheating is that public good non-producers perform poorly in monoculture, but can outcompete producers in mixed culture, where they capitalize on the public goods produced by others^15^. To test these predictions, we first grew the four strains as monocultures in CAA medium supplemented with either iron or a concentration gradient of the synthetic iron chelator 2,2’-bipyridine. We found that the growth of the siderophore non-producer (H111Δ*orbJ*Δ*pchAB*) became significantly compromised compared to the siderophore producers as soon as the iron chelator was added (Fig. S2). Next, we cocultured the siderophore non-producer together with the siderophore producers (wild type, H111Δ*orbJ* or H111Δ*pchAB*) across a range of culturing conditions (Fig. 3). We consistently observed that the non-producer outcompeted the wildtype (*F*_1,120_ = 36.9; *p* < 0.0001) and the pyochelin producer (H111Δ*orbJ*; *F*_1,119_ = 471.3; *p* < 0.0001) under iron-limitation (Fig. 3), and thus acted as a cheater. In stark contrast, the non-producer was unable to cheat on the ornibactin producer (H111Δ*pchAB*) and lost under all conditions (Fig. 3; *F*_1,128_ = 235.5; *p* < 0.0001). In iron-rich media, where siderophores are neither produced nor needed, the fitness differences between the siderophore producers and the non-producer disappeared (Fig. 3a-3c).

**Figure 3.**
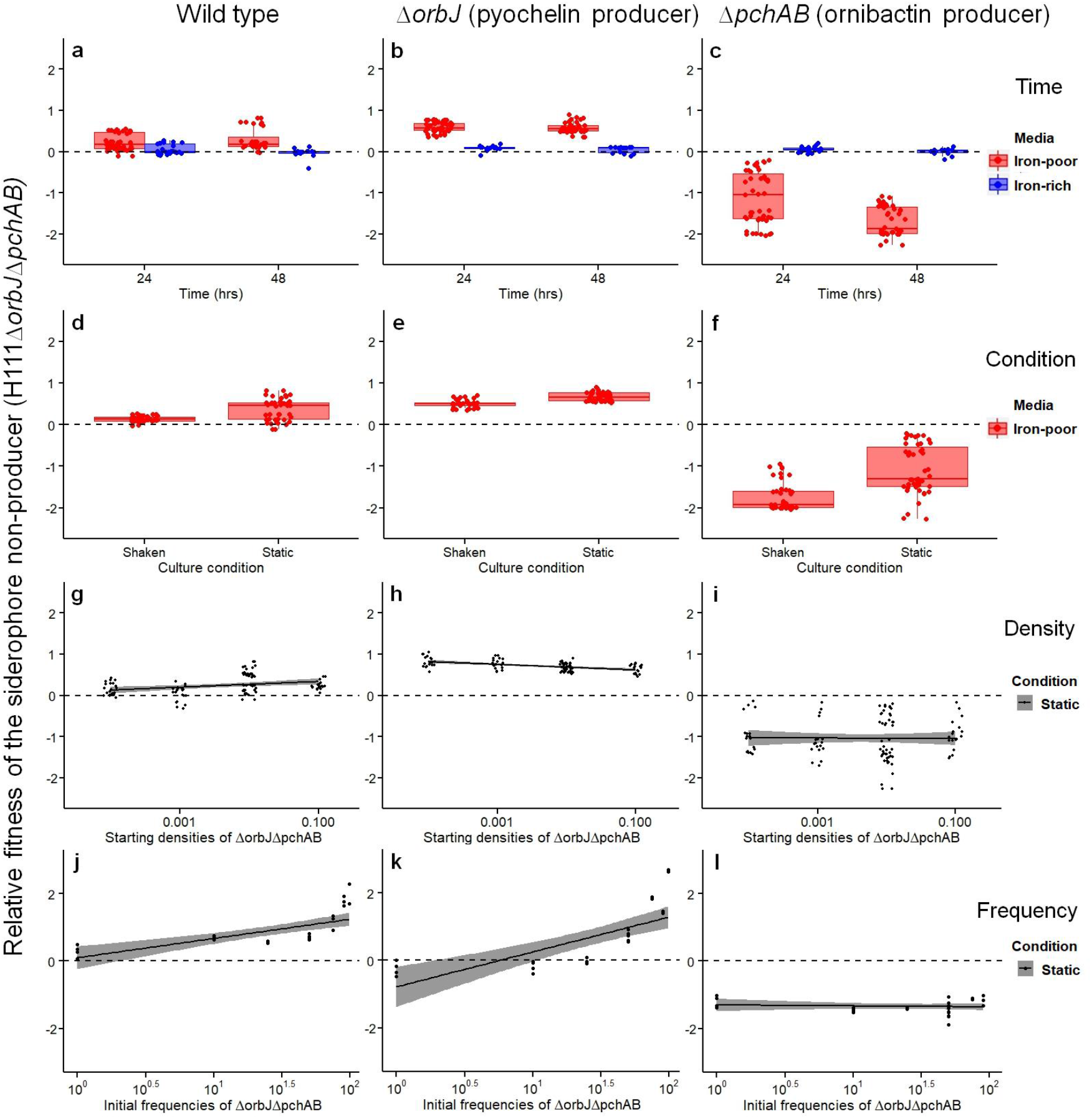
The siderophore non-producer H111Δ*orbJ*Δ*pchAB* can cheat on the wild type and the pyochelin producer, but not on the ornibactin producer. We competed the siderophore non-producer H111Δ*orbJ*Δ*pchAB* tagged with a constitutive mCherry marker against the wild type (first column), the pyochelin producer H111Δ*orbJ* (second column), and the ornibactin producer H111Δ*pchAB* (third column) across a range of culturing conditions. All plots show the relative fitness of the siderophore non-producer, and the dashed horizontal lines represent fitness parity (i.e. when none of the two strains wins the competition). **a-c**, summary of relative fitness values across all conditions, showing that the siderophore non-producer acts as cheater in competition against the wildtype and the pyochelin producer, but loses against the ornibactin producer under iron-poor conditions. Under iron-rich conditions, there is fitness parity between strains, confirming that the observed fitness patterns under iron limitation are mediated by siderophores. **d-f**, relative fitness values of the siderophore non-producer under shaken versus static culturing conditions. **g-i**, relative fitness values of the siderophore non-producer across a range of starting cell densities. **j-l**, relative fitness values of the siderophore non-producer across a range of strain mixing frequencies.

### Culturing conditions affect competition, but often differently as compared to the *P. aeruginosa* pyoverdine system

Work on pyoverdine-sharing in *P. aeruginosa* showed that longer culturing times can lead to more extreme fitness outcomes^28^. We found this to hold true in two of our three competition experiments (Fig. 3a-c). In comparison to an experimental time-course of 24 hrs, the relative fitness of the non-producer after 48 hrs was significantly higher in competition with the wild type (Fig. 3a; *F*_1,42_ = 4.57; *p* = 0.0382), unchanged in competition with the pyochelin producer H111Δ*orbJ* (Fig. 3b; *F*_1,41_ = 0.002; *p* = 0.9578), and significantly lower in competition with the ornibactin producer H111Δ*pchAB* (Fig. 3c; *F*_1,45_ = 36.58; *p* < 0.0001).

Shaking is supposed to increase the mixing of cells and public goods, and has been shown to improve pyoverdine cheating abilities in *P. aeruginosa*^34^ We found the opposite to be the case for the *Burkholderia* siderophores (Fig. 3d-f). Relative fitness of the non-producer was consistently higher in static compared to shaken cultures (Fig. 3d, against wild type: *F*_3,78_ = 12.65; *p* < 0.0001; Fig. 3e, against H111Δ*orbJ*: *F*_3,85_ = 21.71; *p* < 0.0001; Fig. 3f; against H111Δ*pchAB*: *F*_3,86_ = 37.38; *p* < 0.0001).

Previous work indicated that non-producers can exploit siderophores under static conditions more efficiently at high cell density^21,27^. We found that cell density at inoculation had no or only a minor effect on the relative fitness of the non-producer (Fig. 3g, against wild type: *t*_100_ = 1.04; *p* = 0.3021; Fig. 3h, against H111Δ*orbJ*: *t*_97_ = -3.02, *p* = 0.0031; Fig. 3i, against H111Δ*pchAB*: *t*_100_ = -0.518, *p* = 0.6062).

Finally, it was reported that *P. aeruginosa* pyoverdine non-producers were more successful at outcompeting producers when rare^27^. When probing for this relationship in our *Burkholderia* system, we observed the opposite pattern in two out of three cases: non-producers experienced significantly higher relative fitness advantages when more common in competition against the wildtype (Fig. 3j; *F*_1,30_ = 46.2, *p* < 0.0001) and the pyochelin producer H111Δ*orbJ* (Fig. 3k; *F*_1,29_ = 226.6, *p* < 0.0001). In the latter case, the non-producer even lost the competition at initial frequencies smaller than 10%. In competition with the ornibactin producer H111Δ*pchAB*, the non-producer had severe fitness disadvantages at all the frequencies (Fig. 3l; *F*_1,29_ = 1.24, *p* = 0.2730).

### Deletion of the *orbJ* ornibactin synthesis gene reduces mRNA levels of the down-stream receptor gene

Because the ornibactin synthesis gene *orbJ* and the receptor encoding gene *orbA* are part of the same operon, we hypothesize that mutations in *orbJ* might have negative effects on the downstream *orbA* gene (Fig. S3). Such negative effects could include lower *orbA* expression levels or altered mRNA stability, which would in turn curb receptor synthesis and cheating opportunities. To test this hypothesis, we compared the mRNA levels of *orbA* in the wild type relative to our three deletion mutants (Fig. 4a). Compatible with our hypothesis, we found that the amount of *orbA* mRNA was significantly reduced in H111Δ*orbJ* and H111Δ*orbJ*Δ*pchAB*, the two strains lacking the upstream *orbJ* gene (one-tailed t-tests relative to the wild type; H111Δ*orbJ*: *t*_8_ = -21.38, *p* < 0.0001; H111Δ*orbJ*Δ*pchAB*: *t*_8_ = -19.62, *p* < 0.0001). Conversely, *orbA* mRNA levels remained unchanged in H111Δ*pchAB*, which possesses an intact *orbJ* (*t*_8_ = 1.71, *p* = 0.1250). If our hypothesis is correct then negative effects caused by *orbJ* mutations should only affect downstream but not upstream genes. In support of this, we found that mRNA levels were not negatively affected in the upstream gene *orbI*, and even slightly increased in H111Δ*orbJ*(*t*_8_ = 5.34, *p* < 0.001) and H111Δ*orbJ*Δ*pchAB* (*t*_8_ = 11.63, *p* < 0.0001; Fig. 4b).

**Figure 4.**
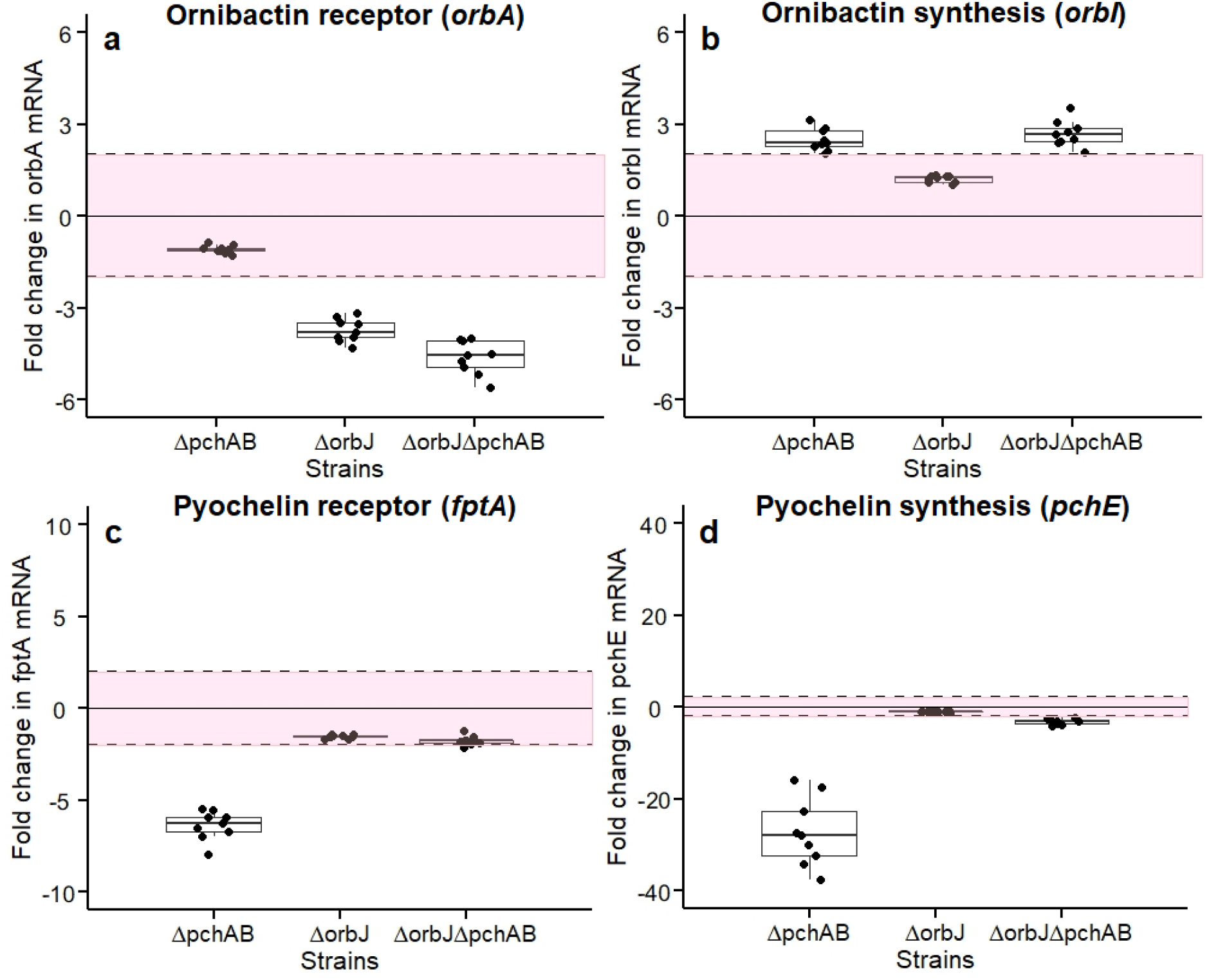
Change in the mRNA levels of siderophore synthesis (*orbI* and *pchE*) and receptor genes (*orbA* and *fptA*) in siderophore mutants relative to the wildtype. For each siderophore system (ornibactin and pyochelin), we used qPCR to quantify changes in mRNA levels in the siderophore synthesis genes *orbI* and *pchE* and the respective receptor genes *orbA* and *fptA* in all siderophore mutants relative to the wildtype. We asked whether the genetic architecture of the siderophore locus affects mRNA levels in mutants lacking the siderophore synthesis genes *orbJ* and/or *pchAB*. a, fold-change in mRNA levels of *orbA*, the ornibactin receptor gene located downstream of the mutated *orbJ* gene. b, fold-change in mRNA levels of *orbI*, an ornibactin synthesis gene located upstream of the mutated *orbJ* gene. c, fold-change in mRNA levels of *fptA*, the pyochelin receptor gene located in an separate operon than the mutated *pchAB* genes. d, fold-change in the expression of *pchE*, a pyochelin synthesis gene located in an separate operon than the mutated *pchAB* genes. All values are scaled relative to the mRNA levels in the wild type. Dashed lines and shaded areas depict the interval [-2|+2], in which mRNA level changes were not considered biologically significant.

We expect such linkage-based negative effects to be absent for pyochelin because the genes for synthesis and uptake are organized in separate operons (Fig. S3)^32^. However, the regulation of pyochelin is complicated for two reasons. First, due to its relatively low iron affinity, pyochelin is typically considered as a secondary siderophore^35^, regulatorily suppressed by primary siderophores such as pyoverdine^36^ and ornibactin^37^. Second, pyochelin itself is a signalling molecule, which triggers its own expression^38^. Thus, it is hard to predict how the deletion of *pchAB* would affect mRNA levels of the *fptA* receptor gene and the pyochelin synthesis gene *pchE* in H111Δ*pchAB* (where signalling is impaired) and in H111Δ*orbJ*Δ*pchAB* (where ornibactin inhibition and signalling are impaired). However, we predicted that mRNA levels for *fptA* and *pchE* should follow the same pattern. In line with this prediction, we observed that the expression of both *fptA* (Fig. 4c) and *pchE* (Fig. 4d) were significantly lower in H111Δ*pchAB* compared to the wildtype (*t*-test; for *fptA*: *t*_8_ = 20.66, *p* < 0.0001; for *pchE*: *t*_8_ = 10.74, *p* < 0.0001), probably due to the impaired signalling. These genes were also similarly expressed in H111Δ*orbJ*Δ*pchAB*, but no longer differentially from the wildtype, probably because ornibactin-mediated repression is released in this mutant.

### Over-expression of *orbA* from plasmids enables H111Δ*orbJ*Δ*pchAB* to cheat on ornibactin producers

To finally prove that the inability of H111Δ*orbJ*Δ*pchAB* to cheat on ornibactin producers is due to reduced receptor availability, we introduced a plasmid with a constitutively expressed *orbA* receptor gene into H111Δ*orbJ*Δ*pchAB*, thereby bypassing the genetic linkage in the operon. We observed that plasmid carriage and/or the overexpression of *orbA* had a metabolic cost, measurable in both monoculture growth assays and in direct competition between H111Δ*orbJ*Δ*pchAB:orbA* and its parental strain H111Δ*orbJ*Δ*pchAB* (Fig. S4). We accounted for these intrinsic fitness differences, which are unlinked to social interactions, in the subsequent two assays.

First, we repeated the supernatant assay shown in Fig. 2, but this time we fed supernatants containing siderophores to the over-expresser H111Δ*orbJ*Δ*pchAB:orbA*. We found that this mutant was now equally well stimulated by the supernatants from the wildtype and H111Δ*pchAB* (producing only ornibactin), suggesting that ornibactin uptake is no longer constrained (Fig. 5a). Our control experiments confirmed again that the growth stimulatory patterns are driven by siderophores, as the observed effects disappeared when H111Δ*orbJ*Δ*pchAB:orbA* was either grown with supernatants from iron-rich media containing little siderophores (Fig. 5b), or with supernatants containing siderophores but replenished with iron (Fig. S5).

**Figure 5.**
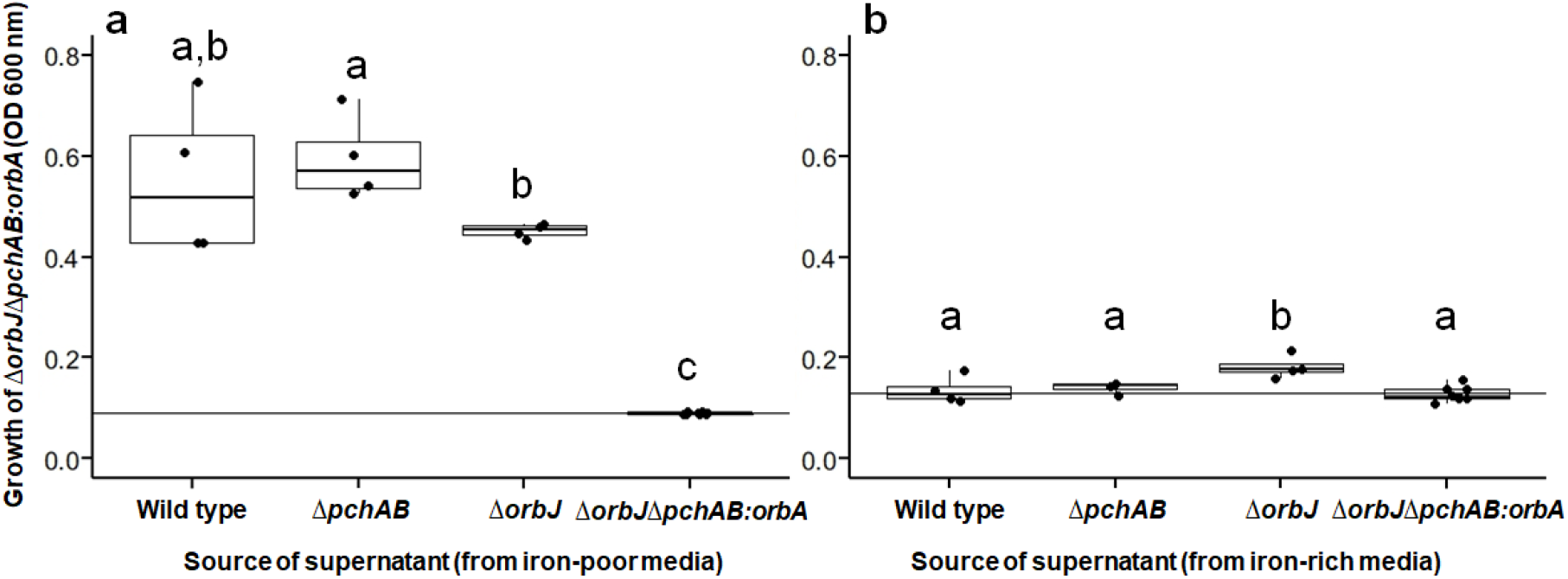
Overexpression of the ornibactin receptor gene from a plasmid restores ornibactin growth stimulation in the siderophore non-producer. We repeated the supernatant assay described in Fig. 2, but this time we supplemented the supernatants from the siderophore producers to H111Δ*orbJ*Δ*pchAB:orbA*, overexpressing the receptor gene *orbA* from a plasmid. a, the supernatant from H111Δ*pchAB*, containing only ornibactin, now stimulated the growth of the siderophore non-producer to a similar levels as the wildtype, and even more so than the supernatant from H111Δ*orbJ*, containing only pyochelin. This demonstrates that the non-producer was no longer short of ornibactin receptors. b, in our control experiments, the growth-stimulatory effects disappeared when supernatants were harvested from iron-rich media that contain little or no siderophores. Different letters above the boxplots indicate statistically significant differences between treatments.

Second, we repeated the competition assays using H111Δ*orbJ*Δ*pchAB:orbA* as the potential cheater strain (Fig. 6). Similar to our previous finding (Fig. 3), we found that this siderophore negative mutant could act as cheater and significantly outcompete the wild type (*t*_23_ = 7.84, *p* < 0.0001) and the pyochelin producer H111Δ*orbJ* (*t*_23_ = 5.93,*p* < 0.0001). However, in stark contrast to Fig. 3, we observed that the *orbA* over-expresser could also successfully cheat and outcompete the ornibactin producer H111Δ*pchAB* (*t*_23_ = 4.80, *p* < 0.0001), indicating that insufficient receptor availability is indeed the cause that constrains cheating in H111Δ*orbJ*Δ*pchAB*.

**Figure 6.**
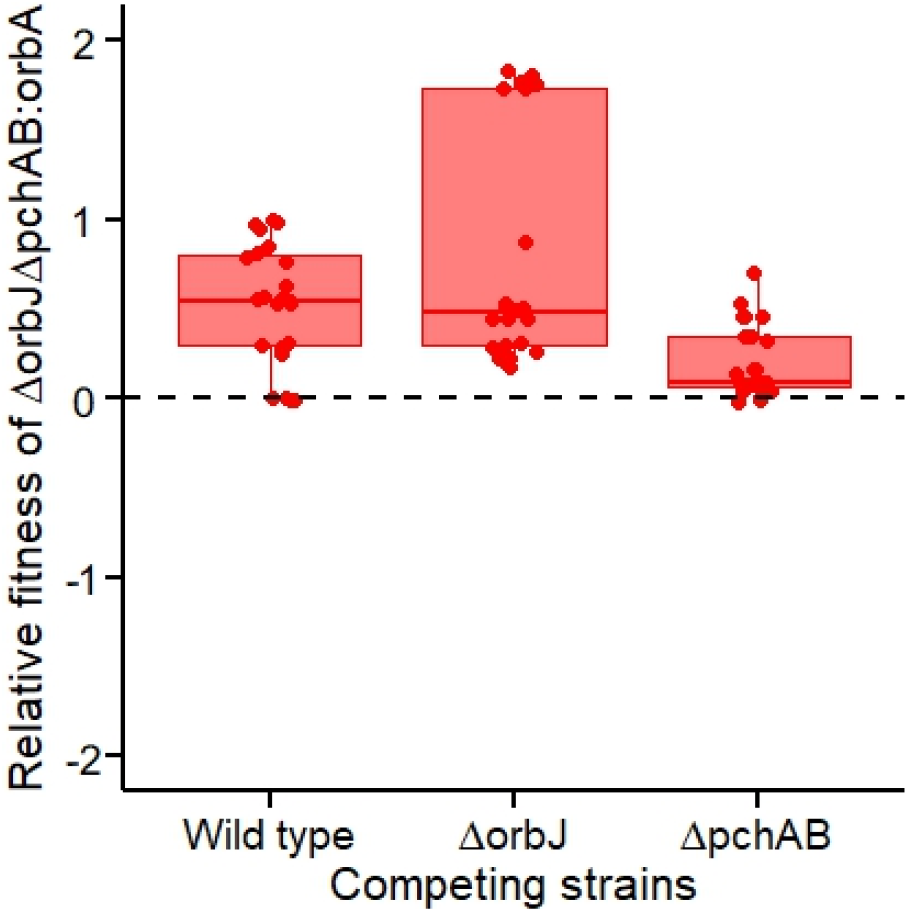
Overexpression of the ornibactin receptor gene from a plasmid allows the siderophore non-producer to efficiently cheat on the wildtype and the ornibactin producer. We repeated the competition experiment described in Fig. 3a-c, but this time we competed the siderophore producers against H111Δ*orbJ*Δ*pchAB:orbA*, overexpressing the receptor gene *orbA* from a plasmid. After a competition period of 24 hours in iron-poor media, we found that this non-producer could significantly outcompete all siderophore producers, including H111Δ*pchAB*, producing only ornibactin (*p* < 0.0001 for all comparisons). The dashed horizontal line represents fitness parity (i.e. when none of the two strains wins the competition). Since the relative fitness values did not significantly differ between static and shaken conditions, the two treatments were merged.

## Discussion

There is a comprehensible tendency in biology to extrapolate from work on model organisms to the large diversity of non-modal systems existing in nature^39^. In this context, research on the production and sharing of pyoverdine by the opportunistic human pathogen *P. aeruginosa* has become a prime example of cooperation in bacteria. Findings from this study system – including the observation that pyoverdine-deficient mutants act as cheaters, potentially bringing cooperation to collapse^7^, and that cheaters perform best in shaken cultures^34^, at high cell density^29^ and when rare^27^ – have often been interpreted as hallmarks of microbial cooperative systems^15,16,40,41^. Here, we studied siderophore production in *B. cenocepacia* H111 and found that patterns of siderophore sharing and exploitation can be very different from the ones reported for *P. aeruginosa*. Most importantly, we observed that secreted ornibactin (the primary siderophore of this species) cannot be exploited by non-producers because the genetic architecture of the ornibactin locus implies that a deficiency in siderophore production leads to the concomitant down-regulation of ornibactin receptor production. Moreover, we found that pyochelin (the secondary siderophore of *B. cenocepacia* H111) can be efficiently exploited by non-producers, but the relative success of cheaters was independent of cell density, and highest in static cultures and when wild-type *B. cenocepacia* was rare, thus opposite to the patterns observed for pyoverdine in *P. aeruginosa*. Our findings highlight that we have to be careful with extrapolations, even for closely-related systems such as pyoverdine and ornibactin, but rather embrace the diversity offered by nature, which can lead, as in our case, to new discoveries and offer a more complete picture of the diversity of social interactions in microbes.

We have discovered a novel mechanism that secures benefits of secreted siderophores to producers and limits the ability of non-producers to partake in public goods use. This mechanism entails tight linkage between the ornibactin synthesis and receptor genes as part of the same operon. A consequence of this linkage is that a deletion in the synthesis gene negatively affects the down-stream receptor gene, such that ornibactin non-producers are short of receptors for ornibactin uptake. Reduced receptor availability is best demonstrated by our supernatant feeding assays showing that the siderophore non-producer H111Δ*orbJ*Δ*pchAB* can still uptake ornibactin to some extent (Fig. 2a), but growth stimulation was much reduced compared to pyochelin and relative to our engineered strain overexpressing the ornibactin receptor gene (Fig. 5a). While we used an engineered strain to demonstrate this effect (in-frame *orbJ* deletion), we suggest that the same effects would arise for many types of natural mutations. Specifically, insertions or deletions that lead to frameshifts, or mutations introducing a stop codon in the up-stream synthesis genes could all detrimentally affect the downstream receptor gene. In support of our argument, Sokol et al.^42^ showed that ornibactin synthesis mutants in *B. cenocepacia* K56-2, created through transposon insertions, were compromised in ornibactin uptake, indicating that insertions lead to the same phenotypes as reported here. Moreover, studies on evolved siderophore non-producers in *P. aeruginosa* revealed that mutations almost exclusively occurred in regulators of siderophore synthesis^11,17^, which in the case of *Burkholderia* may lead to the silencing of the entire operon, including the receptor gene. Only non-synonymous SNPs leading to an amino acid substitution would probably not affect down-stream receptor expression. However, such mutations would most likely also not lead to the abrogation of ornibactin production and thus not turn producers into potential cheaters. These considerations show that the discovered mechanism likely confers a robust way to prevent the spread of ornibactin null mutants as cheaters.

If the linkage of siderophore synthesis and receptor genes within the same operon is such a powerful mechanism to prevent cheating, why is it then not ubiquitous across siderophore systems? For instance, in the case of pyochelin (*B. cenocepacia* and *P. aeruginosa*) and pyoverdine, the regulation of siderophore and receptor synthesis are partly decoupled^31,32,43^. One possible explanation is that there is a trade-off between the prevention of cheating and the flexibility that can be obtained from independent receptor regulation. Flexible receptor regulation could confer an advantage when bacteria arrive in environments where there is already a pool of secreted siderophores available. In this scenario it was shown for *P. aeruginosa* that bacteria rather rely on pyoverdine recycling than on *de novo* production to satisfy their need for iron^44,45^. This economic mechanism, which can only work if cells are able to selectively up-regulate receptor synthesis, also helps siderophore producers to be competitive against cheaters^45^. This indicates that natural selection might offer multiple solutions to cope with cheating, and regulatory linkage is only one of them.

While we have shown that the genetic architecture of a single trait can constrain cheating, other studies have revealed mechanisms against cheating that are based on the regulatory linkage of multiple traits^46–52^. For instance, Dandekar et al.^48^ showed that mutants deficient in the LasIR quorum-sensing (QS) system in *P. aeruginosa* could cheat on cooperative protease production, but exhibited metabolic insufficiencies in certain media, because this QS system also controls the expression of enzymes required for nutrient degradation. While this pleotropic effect prevented cheating in their study system, the view that regulatory linkage between traits has evolved to stabilize cooperation, has recently been challenged^53^. The arguments are that genetic linkage can break and that cooperation likely selects for pleiotropy and not the other way round. Put simply, if a regulatory element like a cooperative QS system is in place, it makes economic sense to recruit more than one trait under this regulon, leading to linkage. We believe that our system is different from the ones considered above, because the synthesis and uptake of siderophores are parts of the same trait, and the genetic architecture thus involves some level of linkage and physical proximity in the genome (Fig. S3)^31,32,43^. However, the fact that linkage can break over evolutionary time scales persists even for our system. Indeed, a comparative analysis on the variation in the ornibactin locus architecture across different *Burkholderia* species reveals that the genetic linkage reported for *B. cenocepacia* is at least partly broken in two species (*B. cepacia* and *B. paludis*) belonging to the same phylogenetic clade, but also in certain species on neighbouring phylogenetic branches (members of the *B. pseudomallei* group, *B. phytofirmans, B. xenovorans)^32^*. Moreover, for one species *(B. paludis)* with broken linkage we know that it can take up ornibactin without producing it^32^. These considerations indicate that the genetic architecture of social traits can evolve^53^, and linkage might only offer a temporary solution to withstand cheating.

We now turn to pyochelin of *B. cenocepacia* and ask why the patterns of successful cheating were so different from the ones reported for pyoverdine in *P. aeruginosa* (Fig. 3). While additional experiments would be required to conclusively answer this question, we can offer the following tentative explanations. Comparative work across more than a hundred bacterial species revealed that siderophores differ fundamentally in their chemical properties, which determine their level of shareability in the community^20^. Pyochelin, for instance, is a lot smaller and more diffusible than pyoverdine and cheaper to produce^35,36^. We propose that these differences might explain some of the variation in the observed fitness patterns (Fig. 3). Because of its larger size, pyoverdine diffuses less readily, and this could explain why the non-producers’ access to this siderophore is increased in shaken cultures and at high cell density. In contrast, the higher diffusivity of pyochelin could lead to a more homogenous distribution of the siderophore across cells even under static and low density conditions, and thus cancel shaking and density effects (Fig. 3e+h). Meanwhile, our observation that pyochelin cheaters perform best when common (Fig. 3k) could be explained by the relatively low pyochelin production costs. The logic is the following: when non-producers are rare, producers can afford losing a low number of cheap molecules without large fitness costs. However, when nonproducers are common, the burden to producers is expected to increase because most pyochelin molecules will be lost to non-producers due to their high diffusivity. The scenario seems different for pyoverdine, where high production costs already impose significant fitness losses at low cheater frequencies, whereas lower diffusivity constrains the cheaters’ access to the siderophore at high frequency^27^.

In summary, we have established a new system to study siderophore-meditated social interactions in bacteria. Our experiments revealed yet unknown dynamics between cooperative producers and exploitative cheaters and identified a novel mechanism of how cooperators can become resistant to cheating. While our study helps to obtain a more nuanced picture on the socio-biology of siderophores, it also highlights that there is likely an enormous diversity of social interactions out there in nature, and by focussing on model organisms such as *P. aeruginosa* we might have so far only looked at the tip of the iceberg.

## Methods

### Bacterial strains

The experiments were performed with *Burkholderia cenocepacia* H111 (LMG 23991), a clinical isolate from a cystic fibrosis patient^54^. This isolate (hereafter referred to as wild type) produces two siderophores, ornibactin and pyochelin^42,55^. This species further possesses a membrane-embedded siderophore-independent iron uptake mechanism^23^, which we did not focus on in this study because its contribution to iron uptake is relatively minor under the culturing conditions used. We further used in-frame deletion mutants defective for either the production of ornibactin (H111Δ*orbJ*), pyochelin (H1111Δ*pchAB*) or both ornibactin and pyochelin (H111Δ*orbJ*Δ*pchAB*); see Mathews et al.^23^ for details on strain construction. For competition assays, we chromosomally tagged the siderophore non-producer strain (H111Δ*orbJ*Δ*pchAB*) with a constitutive fluorescent mCherry marker, inserted at the attTn7 site using a tri-parental mating^56^. The donor and helper strains used for conjugation are listed in the supplementary table 1. We checked whether *mcherry* gene insertion had any adverse effect by comparing the growth of H111Δ*orbJ*Δ*pchAB* and H111Δ*orbJ*Δ*pchAB-mcherry* as monocultures and in competition with each other both in iron-rich and iron-poor media. All the strains were maintained as clonal populations in 25% glycerol stocks at -80°C.

### Media and culturing conditions

Prior to all growth assays, we inoculated strains from -80°C glycerol stocks in 50 ml sterile falcon tubes containing 10 ml Lysogeny broth (LB), and incubated them for approximately 15 hrs at 37°C, shaken at 220 RPM. The overnight cultures were pelleted using a centrifuge (5000 RPM, 22°C, 3 min), aseptically washed and resuspended twice with 0.8% sterile NaCl solution, and adjusted to OD 600nm = 1. For all our experiments, we used casamino acids medium (CAA: 5 g/l casamino acids, 1.18 g/l K2HPO4 × 3H_2_O, 0.25 g/l MgSO4 × 7H2O, 25 mM HEPES). The CAA was supplemented with either 100 μM 2,2’ - bipyridine, a strong iron chelator inducing iron starvation (henceforth, iron-poor), or 100 μM FeCl_3_ to make the medium iron-replete (henceforth, iron-rich). The concentrations of 2,2’ - bipyridine and FeCl_3_ were chosen based on the outcome of an initial growth experiment, where we subjected the wildtype and the mutant strains to a range of 2,2’ -bipyridine and FeCl3 concentrations. This experiment was carried in a 96-well plate, where bacterial strains were inoculated in 200 μl CAA medium at the starting density of OD 600 = 1×10^−4^. The plates were incubated at 37°C in a microplate reader (SpectraMax Plus 384, Molecular Devices, USA) and the growth was monitored by measuring OD at 600 nm every 15 min for 24 hrs. The plates, otherwise maintained as static cultures, were shaken for 10 seconds prior to measuring OD. This preliminary experiment revealed that the growth of H111Δ*orbJ*Δ*pchAB* was significantly compromised with 100 μM 2,2’ -bipyridine, whereas the wildtype and the two single siderophore mutants could still grow well (Fig. S2).

### Siderophore detection by CAS assay

We used the chrome azurol S (CAS) assay^57^ to quantify the amount of siderophores secreted by *B. cenocepacia* into the extra-cellular medium. We grew all strains in 10 ml iron-poor CAA in a shaken incubator at 37°C, 220 RPM. After 24 hrs, the cell cultures were centrifuged (7000 RPM, 5 min, and 22°C), the supernatants collected and filtered through 0.2 μm sterile Whatman filters. We then mixed 0.5 ml of twenty times diluted supernatant with 0.5 ml of freshly prepared CAS reagents. This mixture was incubated in the dark for 30 min and the blue to orange colour change, which is proportional to the amount of siderophores present in supernatant, was measured through absorbance at 630 nm using a spectrophotometer (Ultrospec 2100 pro, Amersham Biosciences, UK). We used the CAA medium as the negative control and calculated the siderophores activity relative to the wild type.

### Supernatant assay

We investigated whether the siderophore non-producing mutant (H111Δ*orbJ*Δ*pchAB*) is able to freely access the ornibactin and pyochelin from the liquid media by growing it in the presence of supernatants generated from donor strains in iron-poor and iron-rich media. For iron-poor conditions, the supernatants from the wild type contains both ornibactin and pyochelin, whereas the supernatants of H111ΔorbJ or H111Δ*pchAB* only contain pyochelin or ornibactin, respectively. For iron-rich conditions and for H111Δ*orbJ*Δ*pchAB*, supernatants should not contain any siderophores. To generate supernatants, we grew the strains in 50 ml falcon tubes either containing 10 ml iron-rich or iron-poor CAA. The tubes were incubated at 37°C with shaking (220 RPM) for 24 hrs, after which we centrifuged the grown cultures at 5000 RPM for 3 min at room temperature, and filter sterilized the supernatants with 0.2 μm filters. The supernatants were either used fresh or stored at -20°C until use. To test whether the siderophore-negative mutant (H111Δ*orbJ*Δ*pchAB*) can benefit from secreted siderophores in the supernatant, we grew this strain in a medium containing 70% CAA (either iron-rich or iron-poor) supplemented with 30% supernatant. We opted for a full-factorial design, where the growth effects of supernatants from both iron-poor and iron-rich conditions on H111Δ*orbJ*Δ*pchAB* were examined in both iron-poor and iron-rich media. We performed the experiments in 96 well plates, with a starting inoculum of OD 600 nm = 0.01, and static incubation at 37°C. We finally measured the growth of the double knockout at OD 600 nm after 24 hrs.

### Competition assays

We conducted competition assays between the siderophore non-producer (H111Δ*orbJ*Δ*pchAB-mcherry*) and the three siderophore producers (wildtype, H111Δ*orbJ*, H111Δ*pchAB*) to investigate whether the siderophore non-producer can act as a cheater, outcompeting the siderophore-producing cooperators in co-cultures. The competition assays were performed in sterile, flat bottom 24 well plates (Falcon) containing 1.5 ml iron-rich or iron-poor CAA. We always grew monocultures alongside with mixed cultures under identical conditions. Since the relative success of a putative cheater strain can vary in response to environmental conditions, we manipulated four important variables in our competition assays: (a) time: we assessed the relative success of competing strains after 24 hrs and 48 hrs; (b) culturing condition: we carried out competition assays under static and shaken (170 RPM) conditions; (c) cell density: mixed cultures were initiated at four different starting densities, OD 600 nm = 1×10^−4^, 1×10^−3^, 1×10^−2^ or 1×10^−1^; (d) strain frequency: strain volumetric mixing ratio was varied from 1:99, 10:90; 25:75, 50:50, 75:25 to 99:1. If not manipulated then the standard culturing condition included a 1:1 strain mix, starting OD 600 nm = 0.01, incubated at 37°C in a static incubator. All the competition experiments were performed in at least 15-fold replication. We used flow cytometer to determine the ratio of the two competitors before and after the competition. Thanks to the mCherry tag H111Δ*orbJ*Δ*pchAB-mcherry* could unambiguously be distinguished from its competitor (Fig. S6). All flow cytometric analyses were conducted on a LSRFortessa flow cytometer (BD Biosciences), where mCherry was excited at 561 nm and fluorescence emission was quantified with a 610/20 nm bandpass filter. We used a relatively low thresholds (200) both for the forward (FSC) and the side scatter (SSC), and adjusted the voltage of the photomultiplier tube (PMT) so that the entire bacterial population was clearly visible on a FSC vs SSC dot plot. The voltage of the PMT was adjusted in a similar fashion to ensure that all mCherry positive cells become visible. For every replicate, 100,000 bacterial events were recorded and both the initial and the final ratio of two strains in a mix was calculated. We used monocultures of H111Δ*orbJ*Δ*pchAB-mcherry* to estimate the frequency of cells that were mCherry negative (e.g. 0.05 % in the case shown in Fig. S6B), and then adjusted the strain frequencies in mixes accordingly. The flow cytometry data were analyzed using the software package FlowJo (TreeStar, USA). Based on the obtained strain ratios we then calculated the relative fitness of the siderophore-deficient strain as: *v* = [*a_t_* x (1 - *a_0_*)] / [*a_0_* x (1 - *a_t_*)], where *a_0_* and *a_t_* are its initial and the final frequencies, respectively^27^. We log-transformed *v*-values, whereby v < 0 indicates a decrease and *v* > 0 an increase in the relative fitness of H111Δ*orbJ*Δ*pchAB-mcherry*.

### Gene expression analysis

We used quantitative real-time PCR (qPCR) to measure mRNA levels of the following five genes (Table S2): (a) *orbI* encoding a non-ribosomal peptide synthetase involved in ornibactin biosynthesis; (b) *orbA* codes for the ferriornibactin receptor; (c) *pchE* encoding a non-ribosomal peptide synthetase involved in pyochelin synthesis; (d) *fptA* codes for the ferripyochelin receptor; and (e) the housekeeping gene *recA* as a control. Fig. S3 shows where these genes are located within their respective siderophore clusters. *B. cenocepacia* strains were grown in 250 ml Erlenmeyer flasks containing 50 ml sterile iron-poor and iron-rich CAA (37°C, shaken) until the cultures reached mid-exponential phase (OD 600 nm = 0.5 in iron-rich media after 6 hrs of inoculation, and OD 600 nm = 0.2-0.5 in iron-poor media after 10 hrs of inoculation). RNA was isolated and purified from three independent cultures as described elsewhere^58,59^. Briefly, 45 ml culture was mixed with 5 ml of 10% cold tris-HCL saturated phenol (pH 8.0) and the cells were harvested using a centrifuge at 4°C, 7000 RPM for 5 min. The cell pellets were immediately flash frozen using a liquid nitrogen and the frozen cell pellets were stored at -80°C until the RNA was isolated. RNA was isolated using a hot acid phenol-chloroform extraction method^58^. The total RNA was purified using RNeasy purification kit (Qiagen) and the contaminating genomic DNA was removed by two rounds of DNase I treatment (Promega I). Once the absence of genomic DNA was confirmed (by using a 40 cycle long PCR and a primer developed to amplify the H111 genomic DNA; see table S2) the RNA was once again purified using RNeasy kit and the quality of RNA was checked by a NanoDrop (ThermoFisher Scientific). First strand cDNA was synthesized using 5μg pure RNA and M-MLV RT (Promega). qPCR was performed using pure cDNA and qPCR kit (Agilent technologies). The expression levels for a given gene in mutants was compared after normalizing its levels to the wild type levels.

### Construction of a strain constitutively expressing the ornibactin receptor (orbA)

To be able to experimentally express the *orbA* receptor gene in the H111Δ*orbJ*Δ*pchAB* background, we cloned an approximately 1kb *orbA* gene fragment into the modified expression plasmid pBBR1MCS5-Tp^60^. The recombinant plasmid, from which *orbA* was constitutively expressed from the lac promoter, was then transformed into *E. coli* TOP 10, and subsequently transferred to H111Δ*orbJ*Δ*pchAB* by conjugation^56^. The recombinant colonies were selected on *Pseudomonas* isolation agar (*Burkholderia* can grow on this medium) supplemented with Trimethoprim (100 μg/ml) and verified by PCR as well as sequencing. We found that the overexpression of *orbA* in H111Δ*orbJ*Δ*pchAB* (H111Δ*orbJ*Δ*pchAB:orbA*) mildly but significantly affected its growth and the relative fitness compared to its parental strain H111Δ*orbJ*Δ*pchAB* both in iron-rich and iron-poor media (Fig. S4). We repeated the supernatant assays and the competition assays described above using H111Δ*orbJ*Δ*pchAB:orbA* to test whether this strain can cheat siderophore producers. For the competition assays we corrected relative fitness values for the intrinsic fitness costs associated with plasmid carriage.

### Statistical analysis

All statistical analyses were performed with R 2.8.0 (http://www.r-project.org). We used Linear Model (LM) to test whether strains produce significantly different amounts of siderophores and whether the growth of H111Δ*orbJ*Δ*pchAB* is affected by the source of the supernatant. For competition assays we used one-sample t-tests to test whether the relative fitness of H111Δ*orbJ*Δ*pchAB* significantly differs from *v* = 0. We further used linear models to test whether the manipulated factors (time, culturing conditions, strain density and frequency) affected the relative fitness of H111Δ*orbJ*Δ*pchAB*. In the cases of multiple comparisons, we adjusted the *p*-values using the Tukey’s HSD method.

## Acknowledgements

We thank Gabriella Pessi, Anne Leinweber and Yilei Liu for their comments on qPCR methods, and the flow cytometry facility of the University of Zurich for technical support. This project has received funding from the European Research Council (ERC) under the European Union’ s Horizon 2020 research and innovation programme (grant agreement n° 681295) to RK. RK was further supported by the Swiss National Science Foundation (PP00P3_165835).

## Author contributions

S.S. and R.K. designed the study. S.S. and A.M. constructed strains and acquired the data, all authors interpreted the data and wrote the paper.

## References

1. Miethke, M. & Marahiel, M. A. Siderophore-based iron acquisition and pathogen control. Microbiol. Mol. Biol. Rev. 71, 413–451, (2007).

2. Hider, R. C. & Kong, X. Chemistry and biology of siderophores. Nat. Prod. Rep. 27, 637–657, (2010).

3. Cassat, J. E. & Skaar, E. P. Iron in infection and immunity. Cell Host Microbe 13, 509–519, (2013).

4. Braud, A., Geoffroy, V., Hoegy, F., Mislin, G. L. A. & Schalk, I. J. Presence of the siderophores pyoverdine and pyochelin in the extracellular medium reduces toxic metal accumulation in *Pseudomonas aeruginosa* and increases bacterial metal tolerance. Environ. Microbiol. Rep. 2, 419–425, (2010).

5. Hesse, E. et al. Ecological selection of siderophore-producing microbial taxa in response to heavy metal contamination. Ecol. Lett. 21, 117–127, (2018).

6. Butaitė, E., Baumgartner, M., Wyder, S. & Kümmerli, R. Siderophore cheating and cheating resistance shape competition for iron in soil and freshwater Pseudomonas communities. Nat. Commun. 8, 414, (2017).

7. Griffin, A., West, S. A. & Buckling, A. Cooperation and competition in pathogenic bacteria. Nature 430, 1024–1027, (2004).

8. Weigert, M. & Kümmerli, R. The physical boundaries of public goods cooperation between surface-attached bacterial cells. Proceedings of the Royal Society B: Biological Sciences 284, 20170631, (2017).

9. Cordero, O. X., Ventouras, L.-A., DeLong, E. F. & Polz, M. F. Public good dynamics drive evolution of iron acquisition strategies in natural bacterioplankton populations. Proc. Natl. Acad. Sci. U.S.A. 109, 20059–20064, (2012).

10. Julou, T. et al. Cell-cell contacts confine public goods diffusion inside *Pseudomonas aeruginosa* clonal microcolonies. Proc. Natl. Acad. Sci. U.S.A. 110, 12577–12582, (2013).

11. Andersen, S. B., Marvig, R. L., Molin, S., Johansen, H. K. & Griffin, A. S. Long-term social dynamics drive loss of function in pathogenic bacteria. Proc. Natl. Acad. Sci. U.S.A. 112, 10756–10761, (2015).

12. Tekwa, E. W., Nguyen, D., Loreau, M. & Gonzalez, A. Defector clustering is linked to cooperation in a pathogenic bacterium. Proceedings of the Royal Society B: Biological Sciences 284, (2017).

13. Vasse, M. et al. Antibiotic stress selects against cooperation in the pathogenic bacterium *Pseudomonas aeruginosa*. Proc. Natl. Acad. Sci. U.S.A. 114, 546–551, (2017).

14. Granato, E. T., Ziegenhain, C., Marvig, R. L. & Kummerli, R. Low spatial structure and selection against secreted virulence factors attenuates pathogenicity in *Pseudomonas aeruginosa*. ISME J. 12, 2907–2918, (2018).

15. West, S. A., Griffin, A. S., Gardner, A. & Diggle, S. P. Social evolution theory for microorganisms. Nat. Rev. Microbiol. 4, 597–607, (2006).

16. Özkaya, Ö., Xavier, K. B., Dionisio, F. & Balbontin, R. Maintenance of microbial cooperation mediated by public goods in single and multiple traits scenarios. J. Bacteriol. 199, e00297–00217, (2017).

17. Kümmerli, R. et al. Co-evolutionary dynamics between public good producers and cheats in the bacterium *Pseudomonas aeruginosa*. J. Evol. Biol. 28, 2264–2274, (2015).

18. O‘Brien, S., Lujan, A. M., Paterson, S., Cant, M. A. & Buckling, A. Adaptation to public goods cheats in *Pseudomonas aeruginosa*. Proc. R. Soc. B 284, 20171089, (2017).

19. Bruce, J. B., Cooper, G. A., Chabas, H., West, S. A. & Griffin, A. S. Cheating and resistance to cheating in natural populations of the bacterium *Pseudomonas fluorescens*. Evolution 71, 2484–2495, (2017).

20. Kümmerli, R., Schiessl, K. T., Waldvogel, T., McNeill, K. & Ackermann. Habitat structure and the evolution of diffusible siderophores in bacteria. Ecol. Lett. 17, 1536–1544, (2014).

21. Scholz, R. L. & Greenberg, E. P. Sociality in *Escherichia coli*: Enterochelin is a private good at low cell density and can be shared at high cell density. J. Bacteriol. 197, 2122–2128, (2015).

22. Agnoli, K., Lowe, C. A., Farmer, K. L., Husnain, S. I. & Thomas, M. S. The ornibactin biosynthesis and transport genes of *Burkholderia cenocepacia* are regulated by an extracytoplasmic function factor which is a part of the fur regulon. J. Bacteriol. 188, 3631–3644, (2006).

23. Mathew, A., Eberl, L. & Carlier, A. L. A novel siderophore-independent strategy of iron uptake in the genus *Burkholderia*. Mol. Microbiol. 91, 805–820, (2014).

24. Mathew, A., Jenul, C., Carlier, A. L. & Eberl, L. The role of siderophores in metal homeostasis of members of the genus Burkholderia. Environ. Microbiol. Rep. 8, 103–109, (2016).

25. Coenye, T. & Vandamme, P. Diversity and significance of *Burkholderia* species occupying diverse ecological niches. Environ. Microbiol. 5, 719–729, (2003).

26. Sousa, S. A., Ramos, C. G. & Leitao, J. H. *Burkholderia cepacia* complex: emerging multihost pathogens equipped with a wide range of virulence factors and determinants. International Journal of Microbiology 2011, (2011).

27. Ross-Gillespie, A., Gardner, A., West, S. A. & Griffin, A. S. Frequency dependence and cooperation: theory and a test with bacteria. Am. Nat. 170, 331–342, (2007).

28. Kümmerli, R., Griffin, A. S., West, S. A., Buckling, A. & Harrison, F. Viscous medium promotes cooperation in the pathogenic bacterium *Pseudomonas aeruginosa*. Proc. R. Soc. B 276, 3531–3538, (2009).

29. Ross-Gillespie, A., Gardner, A., Buckling, A., West, S. A. & Griffin, A. S. Density dependence and cooperation: theory and a test with bacteria. Evolution 63, 2315–2325, (2009).

30. Zhang, X. X. & Rainey, P. B. Exploring the sociobiology of pyoverdin-producing *Pseudomonas*. Evolution 67, 3161–3174, (2013).

31. Visca, P., Imperi, F. & Lamont, I. L. Pyoverdine siderophores: from biogenesis to biosignificance. Trends Microbiol. 15, 22–30, (2007).

32. Butt, A. T. & Thomas, M. S. Iron acquisition mechanisms and their role in the virulence of *Burkholderia* species. Front. Cell. Infect. Microbiol. 7, 460, (2017).

33. Thomas, M. S. Iron acquisition mechanisms of the *Burkholderia cepacia* complex. BioMetals 20, 431–452, (2007).

34. Leinweber, A., Fredrik Inglis, R. & Kümmerli, R. Cheating fosters species co-existence in well-mixed bacterial communities. ISME J. 11, 1179–1188, (2017).

35. Cornelis, P. Iron uptake and metabolism in pseudomonads. Appl. Microbiol. Biotechnol. 86, 1637–1645, (2010).

36. Dumas, Z., Ross-Gillespie, A. & Kümmerli, R. Switching between apparently redundant iron-uptake mechanisms benefits bacteria in changeable environments Proc. R. Soc. B 280, 20131055, (2013).

37. Tyrrell, J. et al. Investigation of the multifaceted iron acquisition strategies of Burkholderia cenocepacia. BioMetals 28, 367–380, (2015).

38. Michel, L., Bachelard, A. & Reimmann, C. Ferripyochelin uptake genes are involved in pyochelin-mediated signalling in *Pseudomonas aeruginosa*. Microbiology 153, 1508–1518, (2007).

39. Levy, A. & Currie, A. Model organisms are not (theoretical) models. The British Journal for the Philosophy of Science 66, 327–348, (2014).

40. Zhou, L., Slamti, L., Nielsen-LeRoux, C., Lereclus, D. & Raymond, B. The social biology of quorum sensing in a naturalistic host pathogen system. Curr. Biol. 24, 2417–2422, (2014).

41. Bruger, E. & Waters, C. Sharing the sandbox: Evolutionary mechanisms that maintain bacterial cooperation. F1000Res. 4, 1504, (2015).

42. Sokol, P. A., Darling, P., Woods, D. E., Mahenthiralingam, E. & Kooi, C. Role of ornibactin biosynthesis in the virulence of Burkholderia cepacia: characterization of pvdA, the gene encoding L-ornithine N^5^-oxygenase. Infect. Immun. 67, 4443–4455, (1999).

43. Youard, Z. A., Wenner, N. & Reimmann, C. Iron acquisition with the natural siderophore enantiomers pyochelin and enantio-pyochelin in *Pseudomonas* species. Biometals 24, 513–522, (2011).

44. Imperi, F., Tiburzi, F. & Visca, P. Molecular basis of pyoverdine siderophore recycling in *Pseudomonas aeruginosa*. Proc. Natl. Acad. Sci. U.S.A. 106, 20440–20445, (2009).

45. Kümmerli, R. & Brown, S. P. Molecular and regulatory properties of a public good shape the evolution of cooperation. Proc. Natl. Acad. Sci. U.S.A. 107, 18921–18926, (2010).

46. Foster, K. R., Shaulsky, G., Strassmann, J. E., Queller, D. C. & Thompson, C. R. L. Pleiotropy as a mechanism to stabilize cooperation. Nature 431, 693–696, (2004).

47. Jousset, A. et al. Predators promote defence of rhizosphere bacterial populations by selective feeding on non-toxic cheaters. ISME J. 3, 666–674, (2009).

48. Dandekar, A. A., Chugani, S. & Greenberg, E. P. Bacterial quorum sensing and metabolic incentives to cooperate. Science 338, 264–266, (2012).

49. Ross-Gillespie, A., Dumas, Z. & Kümmerli, R. Evolutionary dynamics of interlinked public goods traits: an experimental study of siderophore production in *Pseudomonas aeruginosa*. J. Evol. Biol. 28, 29–39, (2015).

50. Wang, M., Schaefer, A. L., Dandekar, A. A. & Greenberg, E. P. Quorum sensing and policing of *Pseudomonas aeruginosa* social cheaters. Proc. Natl. Acad. Sci. U.S.A. 112, 201500704, (2015).

51. Majerczyk, C., Schneider, E. & Greenberg, E. P. Quorum sensing control of Type VI secretion factors restricts the proliferation of quorum-sensing mutants. eLife 5, e14712, (2016).

52. Özkaya, O., Balbontin, R., Gordo, I. & Xavier, K. B. Cheating on cheaters stabilizes cooperation in *Pseudomonas aeruginosa*. Curr. Biol. 28, 2070–2080, (2018).

53. Dos Santos, M., Ghoul, M. & West, S. A. Pleiotropy, cooperation, and the social evolution of genetic architecture. PLoS Biol. 16, e2006671, (2018).

54. Gotschlick, A. et al. Synthesis of multiple N-acylhomoserine lactones is wide-spread among the members of the *Burkholderia cepacia* complex. Syst. Appl. Microbiol. 24, 1–14, (2001).

55. Darling, P., Chan, M., Cox, A. D. & Sokol, P. A. Siderophore production by cystic fibrosis isolates of *Burkholderia cepacia*. Infect. Immun. 66, 874–877, (1998).

56. Choi, K.-H. & Schweizer, H. P. mini-Tn7 insertion in bacteria with single attTn7 sites: example *Pseudomonas aeruginosa*. Nat. Protoc. 1, 153–161, (2006).

57. Schwyn, B. & Neilands, J. B. Universal chemical assay for the detection and determination of siderophores. Anal. Biochem. 160, 47–56, (1987).

58. Pessi, G. et al. Genome-wide transcript analysis of *Bradyrhizobium japonicum* bacteroids in soybean root nodules. Mol. Plant Microbe Interact. 20, 1353–1363, (2007).

59. Lardi, M. et al. σ 54-Dependent Response to Nitrogen Limitation and Virulence in Burkholderia cenocepacia Strain H111. Appl. Environ. Microbiol. 81, 4077–4089, (2015).

60. Kovach, M. E. et al. Four new derivatives of the broad-host-range cloning vector pBBR1MCS,carrying different antibiotic-resistance cassettes. Gene, 175–176, (1995).

