## Supplementary figures for "Genetic architecture constrains exploitation of siderophore cooperation in *Burkholderia cenocepacia*"

### Supplementary Information

This file contains:

- 2 supplementary tables
- 6 supplementary figures
- supplementary references

### Supplementary tables

**Table S1. The plasmid donor strains used in conjugations for tagging H111Δ*orbJ*Δ*pchAB* with *mcherry*.**

| Strain | Phenotype | Source |
|---|---|---|
| *E. coli* S17-1 λpir miniTn7-Ptac-mCherry | The strain carries conjugation elements and mini-Tn7 plasmid with *mcherry* under constitutive promoter. | Rolf Kümmerli's strain collection maintained at the University of Zurich, Switzerland. |
| *E. coli* S17-1 λ pir pUX-BF13 | The strain carries conjugation elements and the conjugation helper plasmid. | |

**Table S2. The qPCR primers used in the study.**

| Primer | Sequence (5´-3´) | Source |
|---|---|---|
| Ornibactin synthesis (*orbI*-forward) | TGAATCTGCGGCTCGACAC | Microsynth, Switzerland |
| Ornibactin synthesis (*orbI*-reverse) | CAGTGTGCGGCGATGTGATA | |
| Ornibactin receptor (*orbA*-forward) | ACTACAGCCGCTTCGACATC | |
| Ornibactin receptor (*orbA*-reverse) | CATCGTGACGGGCGTATACA | |
| Pyochelin synthesis (*pchE*-forward) | GAGTTGCCACGCGTTTCTC | |
| Pyochelin synthesis (*pchE*-reverse) | CATTCGTGCGAGCGGAATG | |
| Pyochelin receptor (*fptA*-forward) | CGATCTACAGCGTCACGGAA | |
| Pyochelin receptor (*fptA*-reverse) | CGCTGACCTTTGCTTTCCAG | |
| Housekeeping gene (*recA*-forward) | GCGATCTTCGACATCCTGTA | |
| Housekeeping gene (*recA*-reverse) | TTCTCGCCGTTGTAGCTGTA | |
| H111 genomic DNA (*ntrC*-forward) | ACAAGGCGGTCGAGTTGAT | |
| H111 genomic DNA (*ntrC*-reverse) | ATAGAACTGCCCGTCCGACA | |

### Supplementary figures

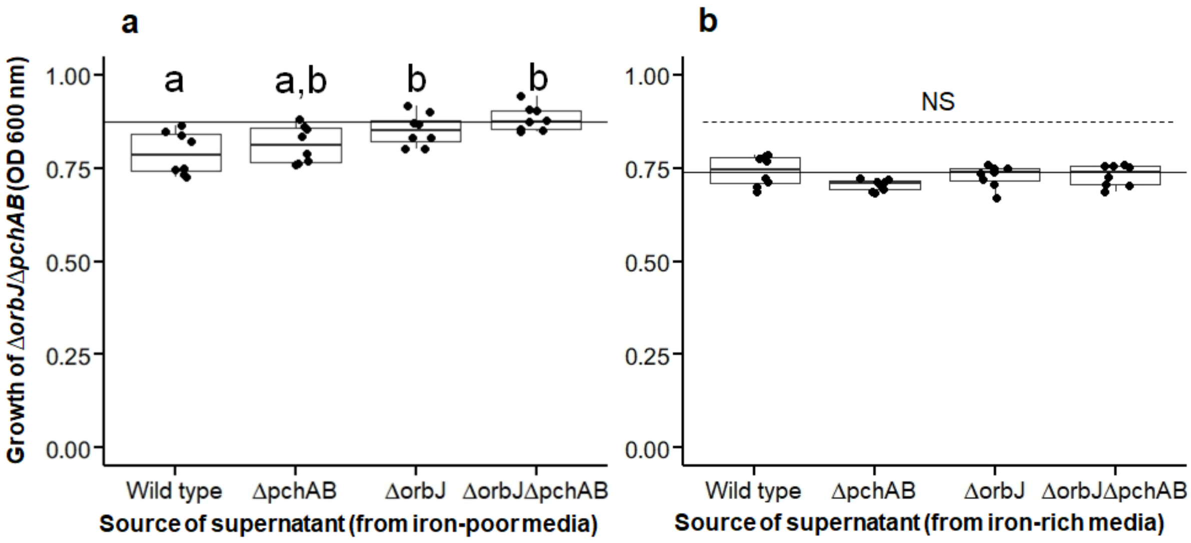


**Figure S1. Control experiments feeding supernatants from siderophore producers to the non-producer H111Δ*orbJ*Δ*pchAB* in iron rich medium.** The siderophore non-producer H111Δ*orbJ*Δ*pchAB* was grown in iron-rich medium for 24 hours, supplemented with supernatants collected from all four strains used in our study (H111 wildtype, H111Δ*pchAB*, H111Δ*orbJ*, H111Δ*orbJ*Δ*pchAB*). **a**, when supernatants from iron-poor medium were supplemented H111Δ*orbJ*Δ*pchAB* grew slightly better in its own supernatant and the supernatant of H111Δ*orbJ* than in the wild type supernatant ($F_{3,28}$ = 5.98; $p$ = 0.0027). **b**, when supernatants from iron-rich medium were supplemented, there was no significant difference between treatments ($F_{3, 28}$ = 2.22; $p$ = 0.1078). Different letters above the boxplots indicate statistically significant differences between treatments.

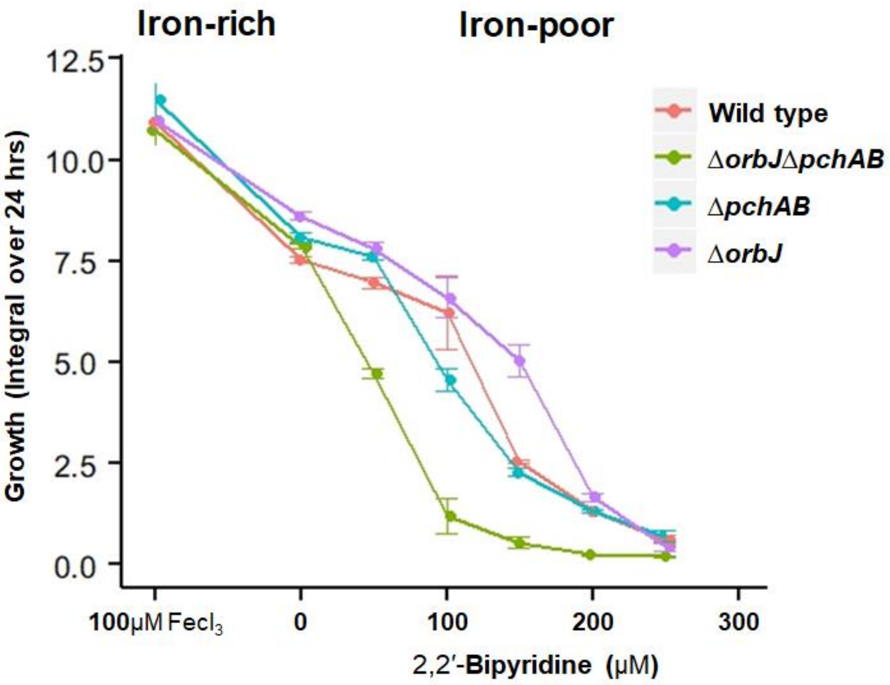


**Figure S2. Growth of *B. cenocepacia* strains across a range of iron availabilities.** *B. cenocepacia* wild type H111 and the three siderophore mutants H111Δ*orbJ*Δ*pchAB* (producing no siderophores), H111Δ*pchAB* (producing ornibactin), and H111Δ*orbJ* (producing pyochelin) were either grown in iron-rich CAA medium supplemented with 100 µM $FeCl_3$ or in iron-poor CAA media (varying 2, 2'-Bipyridine concentration from 50, 100, 150, 200 to 250 µM). All the strains were grown at 37°C and OD 600 nm was monitored every 15 min for 24 hours. The growth curves were analysed in R using the *grofit* package[1]. Because growth trajectories differed fundamentally across conditions, we used spline curve fits and extracted the integral (area under the curve) as growth parameter for comparison. All the strains grew equally well in CAA medium supplemented with 100 µM $FeCl_3$ (iron-rich). Iron depletion (upon the addition of bipyridine) significantly reduced growth of all strains, but most severely for the siderophore non-producer (H111Δ*orbJ*Δ*pchAB*). These results show that siderophores are important for growth.

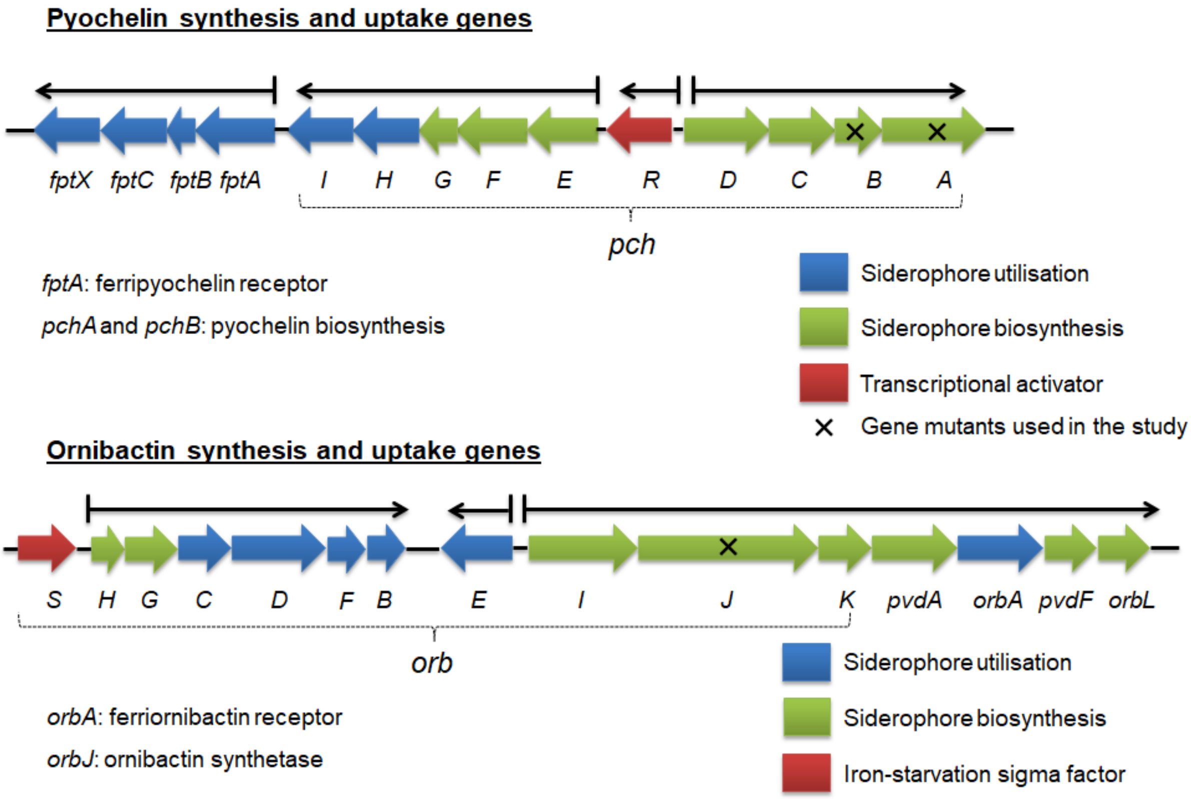


**Figure S3. Genetic architecture of pyochelin and ornibactin genes. a**, the pyochelin synthesis and utilization genes are organized in three different operons (*pchDCBA*, *pchEFGHI* and *fptABCX*). The ferripyochelin receptor gene (*fptA*) is transcribed from a different operon than the synthesis genes *pchAB* (mutated in the study). **b**, the ornibactin synthesis and utilization genes are also organized in three clusters (*orbIJK-pvdA-orbA-pvdF-orbL*, orbE, *orbHGCDFB*). *orbA* and *orbJ* encodes ferriornibactin receptor and non-ribosomal peptide synthetase, respectively. The details of gene organization and regulation can be found in[2-4].

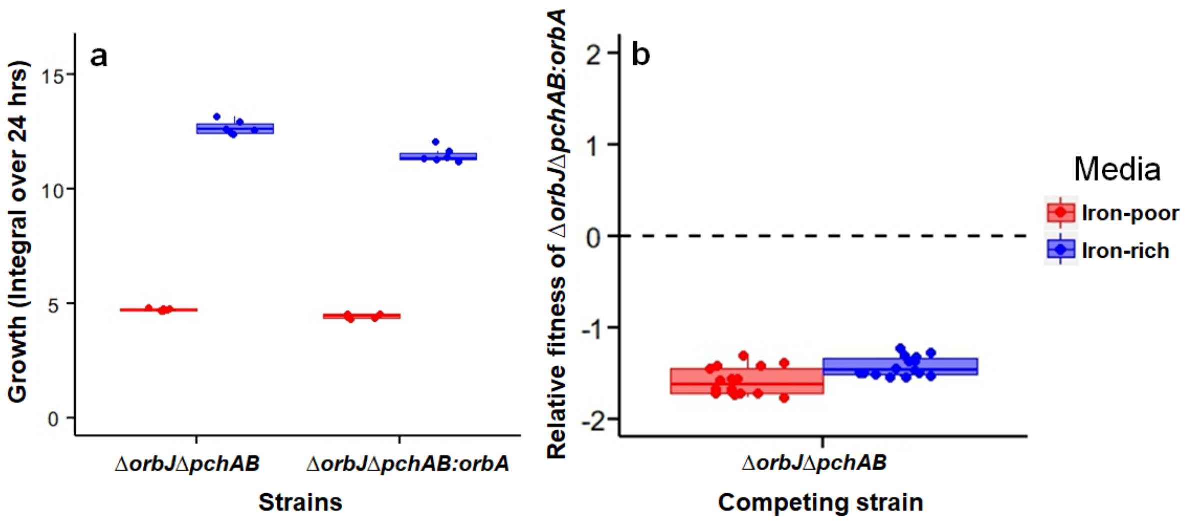


**Figure S4. Monoculture growth and competition between the double mutant (H111Δ*orbJ*Δ*pchAB*) and the *orbA* overexpresser (H111Δ*orbJ*Δ*pchAB:orbA*). a**, the siderophore double mutants H111Δ*orbJ*Δ*pchAB* and H111Δ*orbJ*Δ*pchAB* containing a plasmid from which *orbA* is overexpressed were grown in iron-rich and iron-poor CAA media. Growth at OD 600 nm was monitored every 15 min for 24 hours. We extracted the growth integral and compared it between strains. H111Δ*orbJ*Δ*pchAB:orbA* grew significantly worse than H111Δ*orbJ*Δ*pchAB*, both in iron-rich (*t*-test: $t_{9.9}$, $p < 0.0001$) and iron-poor ($t_{8.2} = 6.49$, $p = 0.0002$) medium. This shows that plasmid carriage has a fitness cost. **b**, costs were confirmed in direct competition assays between the two strains where H111Δ*orbJ*Δ*pchAB:orbA* significantly lost against H111Δ*orbJ*Δ*pchAB* (relative fitness lower than zero) both in iron-rich ($t_{14} = -52.13$, $p < 0.0001$) and iron-poor ($t_{15} = -42.69$, $p < 0.0001$) medium. Competitions were performed under static and shaken conditions, but were combined in the plot because the relative fitness values did not differ between the two conditions.

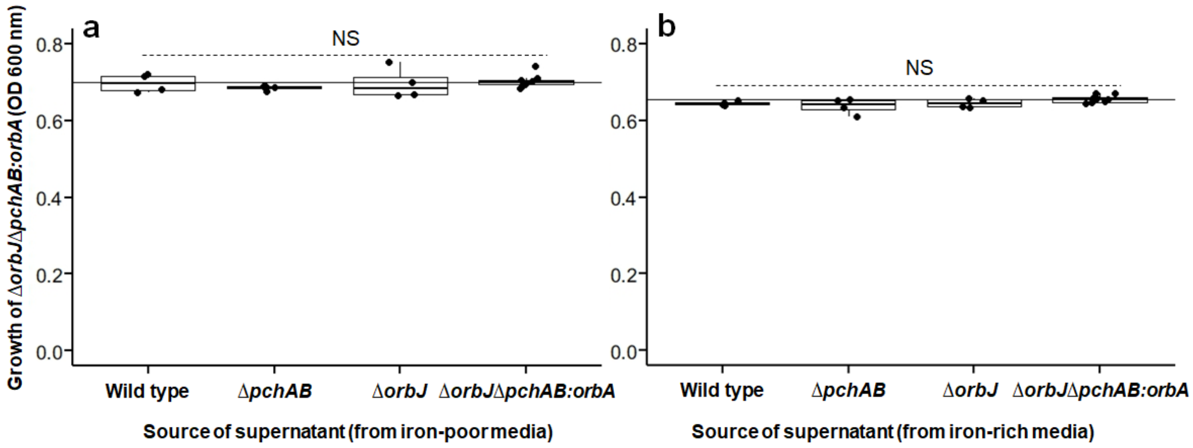


**Figure S5. Control experiments feeding supernatants from siderophore producers to the non-producer H111Δ*orbJ*Δ*pchAB:orbA* overexpressing the ornibactin receptor gene from a plasmid.** The siderophore non-producer H111Δ*orbJ*Δ*pchAB:orbA* was grown in iron-rich medium for 24 hours, supplemented with supernatants from H111 wildtype, H111Δ*pchAB*, H111Δ*orbJ*, and itself. **a**, when supernatants from iron-poor medium were supplemented, there was no significant difference between treatments ($F_{3,16}$ = 0.56, p = 0.6473). **b**, when supernatants from iron-rich medium were supplemented, there was also no significant difference between treatments ($F_{3,16}$ = 2.32, $p$ = 0.1134).

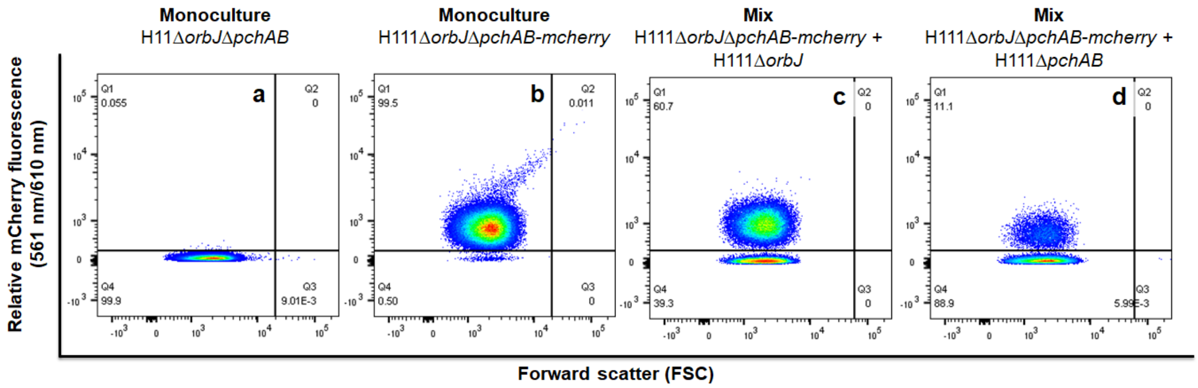


**Figure S6. Examples of flow-cytometry scatter plots from competition experiments between the siderophore non-producer and siderophore producers.** The siderophore double mutant, chromosomally tagged with a constitutively expressed mCherry marker (H111Δ*orbJ*Δ*pchAB-mcherry*) was co-cultured with each of the three siderophore producers (H111 wildtype, H111Δ*pchAB*, H111Δ*orbJ*). With the flow cytometer, we collected approximately 100,000 events from both mono-cultures and mixed cultures before and after a 24-hours competition period. We then plotted the size of cells (forward scatter, FSC) against the mCherry fluorescence to count cells of both types. **a**, a monoculture of the untagged H111Δ*orbJ*Δ*pchAB* strain does not show mCherry fluorescence. This control enabled as to quantify the background fluorescence of cells. **b**, a monoculture of the tagged H111Δ*orbJ*Δ*pchAB-mcherry* strain show relatively strong mCherry fluorescence, with 99.5 % of all cells considered as mCherry positive. **c**, in a 50:50 mix of H111Δ*orbJ*Δ*pchAB-mcherry* and the pyochelin producer (H111Δ*orbJ*) the cells of the two strains can be unambiguously distinguished and their final ratio (60.7:39.3) can be recorded. In this scenario the siderophore non-producer won the competition and acted as a cheater. **d**, the same procedure was applied to a 50:50 mix of H111Δ*orbJ*Δ*pchAB-mcherry* and the ornibactin producer (H111Δ*pchAB*). Here, flow cytometry counts revealed a 11.1:88.9 end ratio, showing that the siderophore non-producer clearly lost the competition and could not act as a cheater.

**Supplementary citations**

1. Kahm, M., Hasenbrink, G., Lichtenberg-Fraté, H., Ludwig, J. & Kschischo, M. grofit: fitting biological growth curves with R. *J. Stat. Softw.* **33**, (2010).
2. Agnoli, K., Lowe, C. A., Farmer, K. L., Husnain, S. I. & Thomas, M. S. The ornibactin biosynthesis and transport genes of *Burkholderia cenocepacia* are regulated by an extracytoplasmic function factor which is a part of the fur regulon. *J. Bacteriol.* **188**, 3631-3644, (2006).
3. Thomas, M. S. Iron acquisition mechanisms of the *Burkholderia cepacia* complex. *BioMetals* **20**, 431-452, (2007).
4. Butt, A. T. & Thomas, M. S. Iron acquisition mechanisms and their role in the virulence of *Burkholderia* species. *Front. Cell. Infect. Microbiol.* **7**, 460, (2017).
